# Segment-specific promoter activity for RNA synthesis in the genome of Oz virus, genus *Thogotovirus*

**DOI:** 10.1101/2024.10.31.621000

**Authors:** Lipi Akter, Ryo Matsumura, Daisuke Kobayashi, Hiromichi Matsugo, Haruhiko Isawa, Yusuke Matsumoto

## Abstract

Oz virus (OZV), a tick-borne, six-segmented negative-strand RNA virus in the genus Thogotovirus, caused a fatal human infection in Japan in 2023. To investigate mechanisms of viral RNA synthesis, we developed an OZV minigenome assay used in mammalian cells. Comparisons of promoter activities across six segments revealed that Segment 5 exhibited markedly lower promoter activity. Unlike the other segments forming a “distal duplex”, a double-strand RNA beginning at the 11th nucleotide on the 5’ end and the 10th on the 3’, Segment 5 partially lacks this feature. Introducing a mutation in Segment 5 distal duplex resulted in a substantial increase in promoter activity. Further, we examined the promoter structures of other viruses in the genus *Thogotovirus* using public database. Thogoto virus Segment 1 also partially lacks a base pair in the distal duplex. In six-segmented RNA viruses, promoter activities are varied, with notable differences in activities likely existing among segments.

**Highlights:**

- A minigenome assay was developed to elucidate the RNA synthesis mechanism of Oz virus.
- Segment 5 of the Oz virus exhibits significantly lower promoter activity compared to other segments.
- The distal duplex region formed by double-strand RNA at the genomic ends is essential for Oz virus RNA synthesis.
- Promoter activities in six-segmented RNA viruses are inherently variable among segments.

## Introduction

The genus *Thogotovirus*, within the family *Orthomyxoviridae*, consists of viruses predominantly transmitted by various species of hard and soft ticks (Hubálek et al., 2014). Oz virus (OZV), a novel member of this genus, was first isolated from a pool of three *Amblyomma testudinarium* nymphs collected in Ehime Prefecture, Japan in 2018 (Ejiri et al., 2018). To date, there has been only one documented case of OZV infection to human reported in Japan. The patient exhibited symptoms of fatigue, loss of appetite, vomiting, joint pain, and fever, and ultimately succumbed to myocarditis. Postmortem examinations, including laboratory tests and pathological assessments, confirmed the diagnosis of viral myocarditis (https://www.niid.go.jp/niid/images/cepr/OZV/230705_OZVfact_eng.pdf).

The genus *Thogotovirus*, which includes OZV, consists of segmented negative-sense single-strand RNA viruses with six segments. OZV is thought to share a genome replication mechanism similar to that of other *Thogotovirus* species. The negative-sense RNA genome is encapsidated by nucleoprotein (NP), allowing it to be recognized as a template by the viral RNA-dependent RNA polymerase (RdRp) (Dick et al., 2024). This RdRp complex is composed of three proteins: PA, PB1, and PB2, and is responsible for both genome replication and mRNA transcription.

The detailed replication mechanism of the OZV genome remains largely unexplored. The minigenome assay of negative-strand RNA viruses is a valuable experimental tool that allows for the analysis of genome replication and mRNA transcription mechanisms mediated by viral RdRp, without the need for using infectious viruses (Hao et al., 2020; Hoenen et al., 2011; Ren et al., 2021; Yamaoka et al., 2022). The minigenome assay enables research aimed at elucidating the molecular basis of viral replication and facilitates experiments for the development of antivirals. In this study, we established a minigenome assay to unravel the replication mechanism of the OZV genome.

## Materials and Methods

### Cells

Human Embryonic Kidney 293 (HEK293) cells and BHK/T7-9 cells (Ito et al., 2003) were cultured in Dulbecco Modified Eagle’s Medium with 10% fetal calf serum and penicillin/streptomycin. Cells were cultured at 37°C in 5% CO_2_.

### Plasmid construction

The Nluc-expressing OZV minigenome plasmid (OZV-Nluc) was constructed using the pCAGGS plasmid backbone. The Nluc gene was flanked by the 3’ and 5’ untranslational regions (UTR) of the OZV EH8 strain genome (GenBank accession numbers: Segment 1; NC_040730.1, Segment 2; NC_040731.1, Segment 3; NC_040732.1, Segment 4; NC_040735.1, Segment 5; NC_040733.1, and Segment 6; NC_040734.1 with some modifications). The minigenome is set under the control of the RNA polymerase-II promoter, and the transcript expressed as a negative sense RNA is cleaved at both ends by a hammerhead ribozyme and a hepatitis delta virus ribozyme (Fig. 1A) (Beaty et al., 2017). The OZV NP, PB1, PB2 and PA genes, which are encoded in Segments 5, 2, 1 and 3 respectively, are set into the multiple cloning site of pCAGGS vector. All OZV plasmids were generated by GenScript’s custom gene synthesis services (GenScript Japan, Tokyo, Japan). A pCAGGS-derivative plasmid for the expression of firefly luciferase (Fluc) was also constructed (Ashida et al., 2024).The mutations in OZV-Nluc were introduced by a standard cloning method.

**Figure 1.**
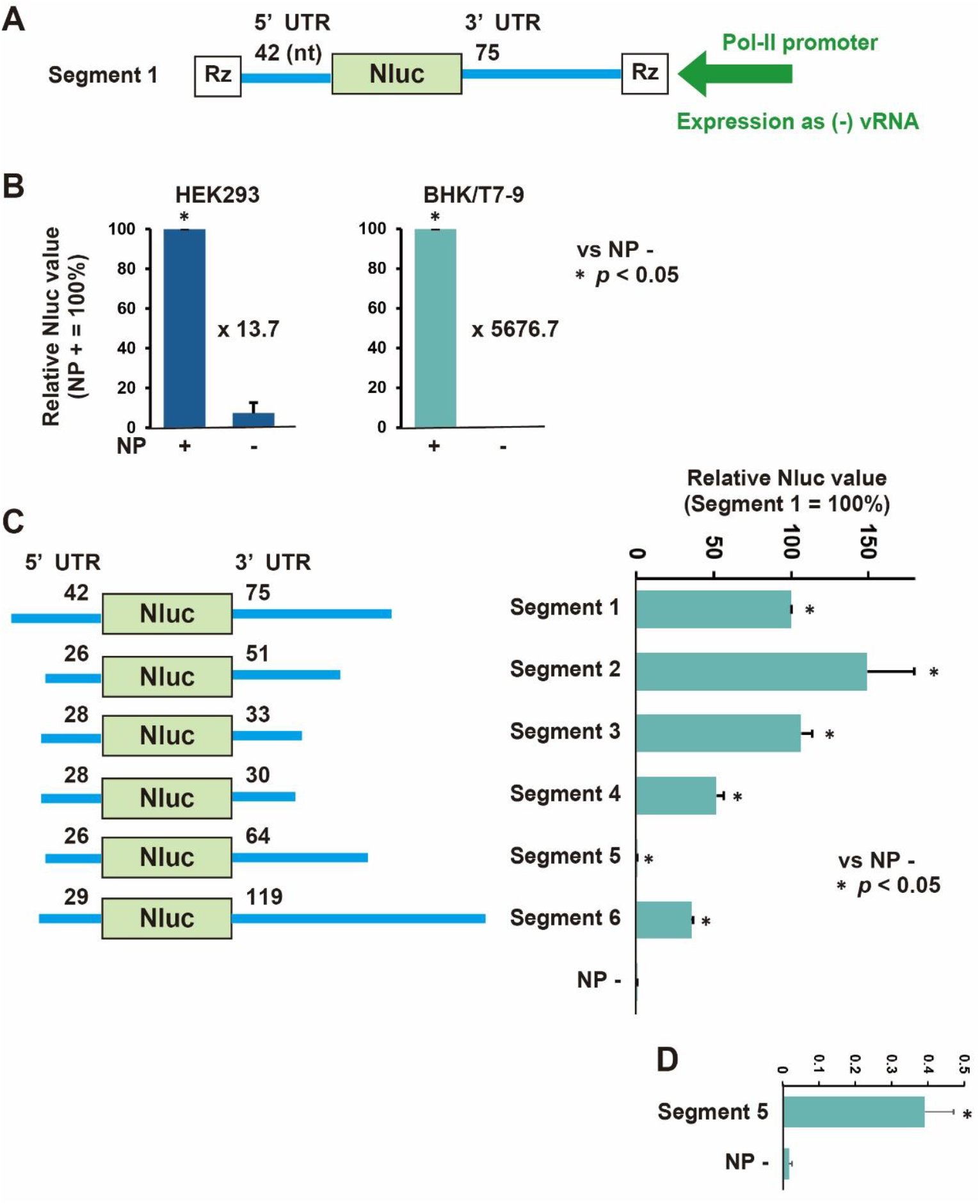
Establishment of OZV minigenome assay. **(A)** A schematic of OZV segment 1 minigenome. The Nluc gene was flanked by the 42 nucleotides (nt) 5’ UTR and 75 nt 3’ UTR of OZV segment 1. The minigenome is set under the control of the RNA polymerase-II promoter, and the transcript expressed as a negative sense (-) viral RNA (vRNA) is cleaved at both ends by ribozymes (Rz). **(B)** Comparison of OZV minigenome activity between HEK293 and BHK/T7-9 cells. Relative Nluc values are shown (NP + equals 100%). **(C)** Comparison of the minigenome activity among OZV segment 1 to 6. The length of 5’ and 3’ UTR of each segment is shown. Relative Nluc values are shown (Segment 1 equals 100%). **(D)** Magnified picture of the results from OZV segment 5. Data represent means and standard deviations from three different experiments. NP – indicates the values using an empty plasmid instead of OZV NP during an experiment using OZV segment 1 minigenome.

### OZV Nluc minigenome assay

The OZV Nluc minigenome assay was performed in HEK293 or BHK/T7-9 cells cultured in 12-well plates (1 x 10^5^ cells/well). Plasmids; OZV-Nluc (0.4 μg), pCAGGS-NP (0.4 μg), -PB1 (0.4 μg), -PB2 (0.4 μg) and -PA (0.4 μg) or empty vector, and Fluc (0.05 μg) were transfected using XtremeGENE HP (Merck, Darmstadt, Germany) according to the manufacturer’s instructions. At 48 hour post-transfection, the cells were lysed with Passive Lysis Buffer and the Nluc and Fluc activities were measured using the Nano-Glo Dual-Luciferase Reporter Assay System (Promega, Madison, WI, USA) according to the manufacturer’s instructions. All activity values measured for Nluc were normalized to the expression levels of Fluc.

### Statistical analysis

A two-tailed unpaired Student’s t-test was employed to compare the means of the two independent groups. The test was performed in R using the t.test() function. The t-statistic and *p*-value were calculated, and a *p*-value of less than 0.05 indicated a statistically significant difference between the groups.

### Results and Discussion

The minigenome derived from OZV Segment 1 has a 5’ UTR of 42 nucleotides and a 3’ UTR of 75 nucleotides (Fig. 1A). This minigenome was transfected into HEK293 or BHK/T7-9 cells along with OZV NP, PA, PB1, and PB2 expression plasmids. As a negative control, an empty vector was used in place of NP (NP-), and a Fluc-expressing plasmid was used as an internal control. At 48 hours posttransfection, Nluc and Fluc expression levels were measured, and the relative Nluc value was compared between the conditions with and without the NP-expressing plasmid (Fig. 1B). In HEK293 cells, the NP+ condition showed 13.7 times higher activity compared to the NP-condition. In contrast, in BHK/T7-9 cells, the NP+ condition demonstrated 5676.7 times higher activity than the NP-condition. Therefore, BHK/T7-9 cells were chosen for subsequent experiments.

Next, we constructed minigenomes derived from all six segments of OZV. As shown in Fig. 1C (left panel), the minigenomes for Segments 1, 2, 3, 4, 5, and 6 had 5’ UTR lengths ranging from 26 to 42 nucleotides, and 3’ UTR lengths ranging from 30 to 119 nucleotides. Using these minigenomes, we investigated the relative Nluc activity, with the activity of the Segment 1 minigenome set at 100% (Fig. 1C). Significant Nluc activity was observed in all segments with NP+ condition compared to the NP-condition. Segment 2 and Segment 3 showed activities of 149% and 106%, respectively. Segment 4 and Segment 6 exhibited activities of 51% and 36%, respectively. However, Segment 5 was notably characterized by extremely low activity, at only 0.4% (Fig. 1D).

In Thogoto virus (THOV), which is closely related to OZV, the 5’ and 3’-genomic ends are crucial for functioning as a promoter. The 1-9th nucleotide of the both 5’ and 3’ ends shows potent complementarity, and the complementarity between the 11-12th nucleotides of the 5’ end and the 10-11th nucleotides of the 3’ end is important for THOV promoter activity (Leahy et al., 1997). Based on this information, we analyzed the complementarity of the 1-9th nucleotides at the 5’ and 3’ ends of all six segments of the OZV genome, as well as the complementarity beyond the 11th nucleotide of the 5’ end and the 10th nucleotide of the 3’ end (Fig. 2A). Prior to this investigation, we revalidated the terminal sequences of the genome using a rapid amplification of cDNA ends (RACE) method and made three corrections to the sequence of Segment 6 in the OZV EH8 strain (NC_040734.1) (Fig. 2A: nucleotides in Segment 6 shown in blue). In all segments, the 1st to 9th nucleotides at both the 5’ and 3’ ends partially formed complementary strands, with 6 to 7 base pairs observed. Additionally, in Segments 1, 2, 3, 4 and 6, complementarity was observed between 6 and 9 base pairs from the 11th nucleotide at the 5’ end and the 10th nucleotide at the 3’ end. In contrast, in Segment 5, a non-complementary region was found between the 12th nucleotide (A) at the 5’ end and the 11th nucleotide (C) at the 3’ end. By changing the 12th nucleotide A at the 5’ end to G, this region is able to form a complementary strand with the 11th nucleotide C at the 3’ end. We prepared a modified Segment 5 minigenome (5’A12G) incorporating this change (Fig. 2B). When the relative minigenome activity was measured with Segment 1 set at 100%, the original Segment 5 showed an activity of 0.4%, whereas Segment 5 (5’A12G) exhibited a significantly higher activity of 252% (Fig. 2B).

**Figure 2.**
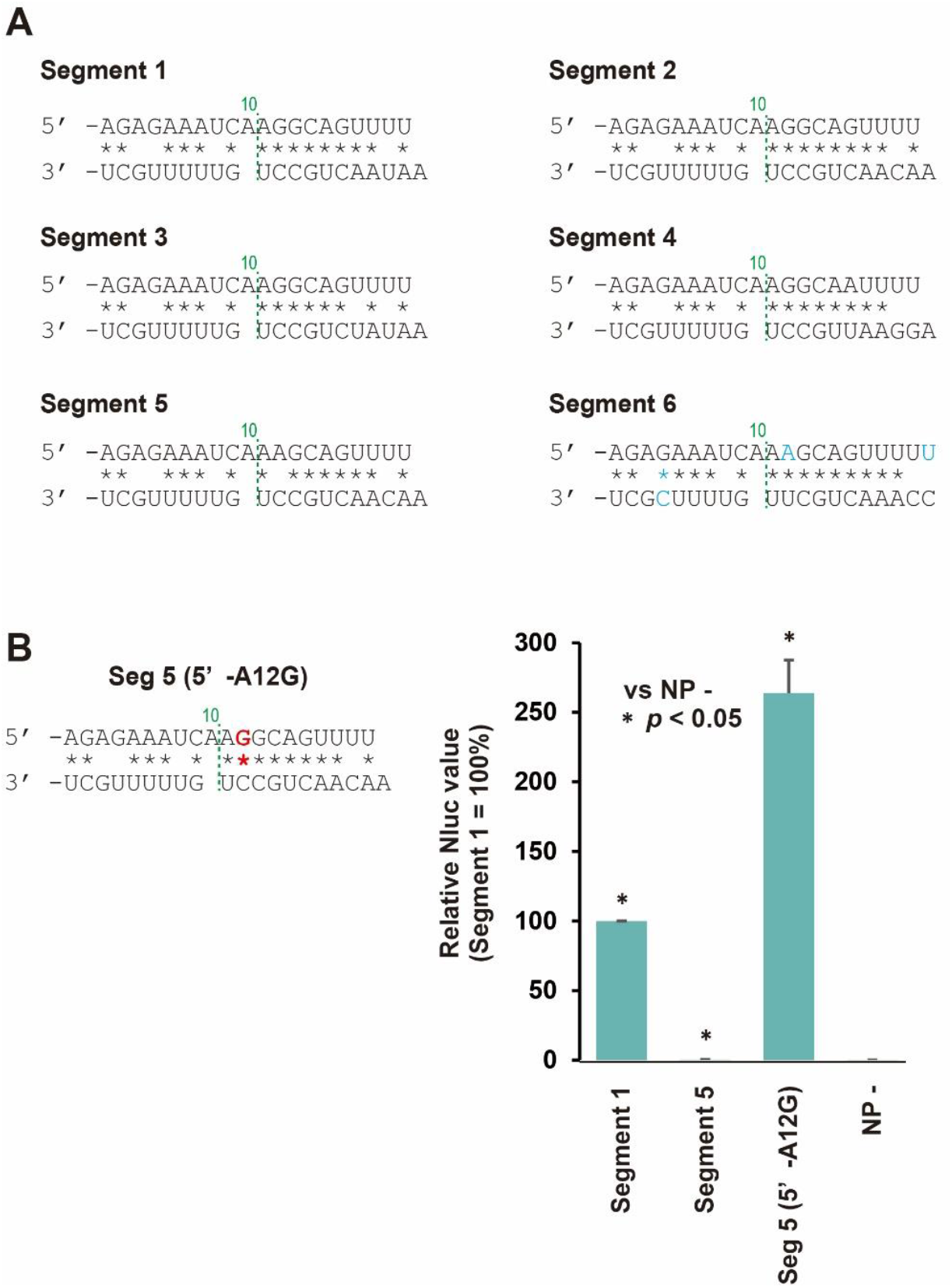
Importance of the base-pairing of 5’-12^th^ and 3’-11^th^ nucleotides for OZV minigenome activity. **(A)** The promoter structure of OZV segments 1-6. The complementarity between the nucleotides at positions 1–9 of the 5’ and 3’-genomic ends, as well as the complementarity from the 11th nucleotide onward at the 5’-genomic end and the 10th nucleotide onward at the 3’-genomic were analyzed. **(B)** The minigenome activity using a mutant minigenome OZV seg 5 (5’-A12G). Relative Nluc values are shown (Segment 1 equals 100%). Data represent means and standard deviations from three different experiments. NP – indicates the values using an empty plasmid instead of OZV NP during an experiment using OZV segment 1 minigenome.

The interaction mode of THOV RdRp with the 5’ and 3’-genomic ends has been analyzed in detail using cryo-electron microscopy (Xue et al., 2024). The nucleotide sequences used for this analysis were derived from THOV Segment 5 (Fig. 3; THOV Segment 5) (Xue et al., 2024). When THOV RdRp forms an RNA synthesis initiation conformation, nucleotides 1-8 at the 3’-genomic end are positioned in the RdRp active site. The nucleotides 1-10 at the 5’-genomic end form a hook structure through intra-strand base pairing in the 5’-hook binding pocket within the RdRp, as seen in other segmented negative-strand RNA viruses (Malet et al., 2023; Pflug et al., 2017). Furthermore, nucleotides 11-17 at the 5’-genomic end and nucleotides 10-16 at the 3’-genomic end form a complementary strand of seven base pairs, resulting in a distal duplex structure. This distal duplex structure is commonly found in other segmented negative-strand RNA viruses although the length of the base pairs is different among viruses (Malet et al., 2023; Neriya et al., 2022; Pflug et al., 2017). Based on this, we proposed a model for the interaction mode of OZV RdRp and the 5’-and 3’-genomic ends (Fig. 3; OZV Segment 1 and 5). It is considered that when nucleotides 1-8 at the 3’-genomic end are positioned in the RdRp active site, the 5’-genomic end also forms a similar hook structure with that of THOV. In Segment 1, it is expected that nucleotides 11-18 at the 5’-genomic end and nucleotides 10-17 at the 3’-genomic end form a distal duplex composed of eight base pairs. This eight base-pairs distal duplex would be also observed in Segments 2 and 4, but in Segment 3, it would form six base pairs, and in Segment 6, nine base pairs would be formed. Considering the minigenome activity of each segment (Fig. 1C), there does not appear to be a strong correlation between the length of the base pairs in the distal duplex and polymerase activity.

**Figure 3.**
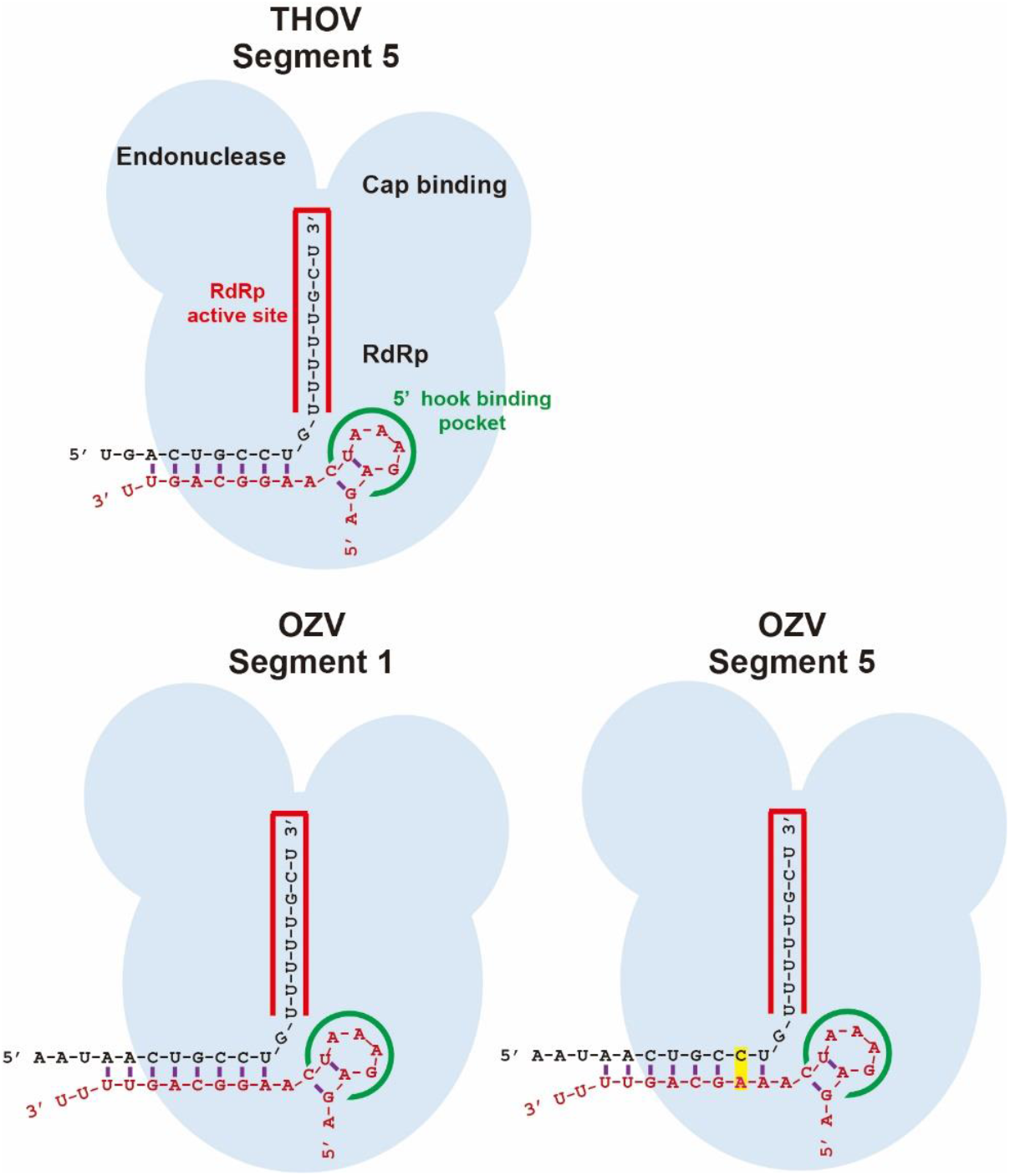
Model of the interaction of genomic ends and RNA-dependent RNA polymerase of THOV and OZV. Schematics of THOV and OZV RdRp composed of PA, PB1 and PB2, and its interaction to 3’- and 5’-genomic ends at an RNA synthesis initiation conformation. The RdRp consists of PA, PB1 and PB2, all of which form an RdRp domain, endonuclease domain and cap-binding domain. An unpaired nucleotides 12^th^ nucleotide from 5’-genomic end and 11^th^ nucleotide from 3’-genomic end in OZV Segment 5 are emphasized in yellow.

In OZV Segment 6, although the 12th nucleotide at the 5’ end differs from the G found in other segments (1, 2, 3, and 4), the 11th nucleotide at the 3’ end is U, allowing the complementarity to be maintained. This slight difference of the nucleotide sequence may be related to the relatively lower activity of Segment 6 (36% of Segment 1) (Fig. 1C).

In OZV Segment 5, which showed particularly low activity, a region in the distal duplex was found where one nucleotide failed to form a complementary strand (Fig. 3; OZV Segment 5). By changing the 12th nucleotide A at the 5’-genomic end to G, allowing the formation of a complementary strand (5’A12G), a significantly high minigenome activity of 252% compared to Segment 1 was observed (Fig. 2B). This indicates that the ability to form six to nine consecutive base pairs in the distal duplex is crucial for RNA synthesis of OZV.

We investigated the promoter structures consisting of the 20 nucleotides from the 5’ and 3’ genomic ends of viruses belonging to the genus *Thogotovirus* registered in the National Center for Biotechnology Information (NCBI) database (Fig. 4). In addition to THOV, we confirmed that the terminal sequences of all six segments are also available for the Bourbon virus, Upolu virus, Aransas virus and Dhori virus (Anderson and Casals, 1973; Briese et al., 2014; Kosoy et al., 2015). We analyzed the complementarity between the nucleotides at positions 1–9 of the 5’ and 3’-genomic ends, as well as the complementarity from the 11th nucleotide onward at the 5’-genomic end and the 10th nucleotide onward at the 3’-genomic end for these sequences (Fig. 4). In all six segments of all viruses, at least five base pairs were observed between nucleotides 1-9 from the 5’ and 3’-genomic ends. Moreover, in all six segments of all viruses, except for THOV Segment 1, six or more consecutive base pairs were observed in the distal duplex region, formed from the 11th nucleotide onward at the 5’-genomic end and the 10th nucleotide onward at the 3’-genomic end. This suggests that the formation of six or more consecutive base pairs in the distal duplex, as seen in OZV, is important for RNA synthesis, and that this mechanism is likely conserved across viruses in the genus *Thogotovirus*.

**Figure 4.**
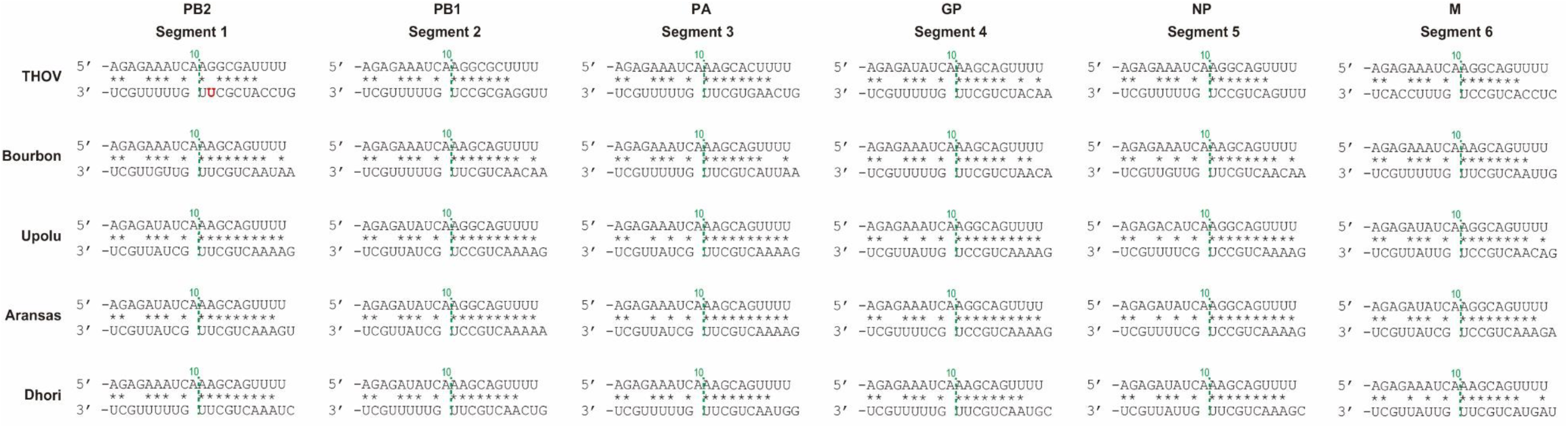
Comparison of the promoter structures of viruses in the genus *Thogotovirus*. The complementarity between the nucleotides at positions 1–9 of the 5’ and 3’-genomic ends, as well as the complementarity from the 11th nucleotide onward at the 5’-genomic end and the 10th nucleotide onward at the 3’-genomic were analyzed. The segment number and encoding genes are shown above. Accession numbers: THOV (NC_006508.1, NC_006495.1, NC_006496.1, MT628407.1, NC_006507.1, NC_006504.1); Bourbon virus (OP594804.1, NC_078584.1, NC_078588.1, NC_078586.1, NC_078587.1, OL989467.1); Upolu virus (NC_078649.1, NC_078651.1, NC_078653.1, NC_078652.1, NC_078654.1, NC_078650.1); Aransas (KC506162.1, KC506163.1, KC506164.1, KC506165.1, KC506166.1, KC506167.1); Dhori virus (NC_034261.1, NC_034263.1, NC_034254.1, NC_034255.1, NC_034262.1, NC_034256.1).

In THOV segment 1, the 12th nucleotide at the 5’ end and the 11th nucleotide at the 3’ end do not form a base pair, which resembles the promoter structure pattern seen in OZV segment 5. Therefore, it is likely that THOV segment 1 exhibits lower promoter activity compared to segments 2–6. While viruses of the genus *Thogotovirus* have six genome segments, the promoter activity does not seem to be uniform across all segments. The observation of extremely low activity in certain segments, such as OZV segment 5, suggests that factors other than the nucleotide sequence at the genomic ends may compensate for this low activity. Alternatively, the presence of segments with low activity may play a role in regulating the replication levels of each segment, which is necessary to maintain a balanced viral life cycle in infected cells.

## Funding

This work was supported by grants from the Japan Agency for Medical Research and Development (AMED) Research Program on Emerging and Re-emerging Infectious Diseases JP23fk0108687 (to Y.M.) and JP22fk0108625 (to H.I.), the JSPS KAKENHI Grant Number 24K09229 (to Y.M.), the Takeda Science Foundation (to Y.M.), the Kato Memorial Bioscience Foundation (to Y.M.) and the Kieikai Research Foundation (to Y.M.). This work was supported by the Cooperative Research Program of Institute for Life and Medical Sciences, Kyoto University, and the Grant for International Joint Research Project of the Institute of Medical Science, the University of Tokyo.

## Acknowledgements

We thank Dr. Naoto Ito (Gifu University) for providing the BHK/T7-9 cells. We also thank Dr. Kyoko Tsukiyama-Kohara (Kagoshima University) and members of her laboratory for their support with the biological experiments.

## Competing interests

Yusuke Matsumoto receives compensation from Denka Co., Ltd. The other authors declare no competing interests.

